# A Comparative assessment of T-Cell response of Healthy donors and Acute Graft-versus-Host-Disease Patients: Customizing immune monitoring platform

**DOI:** 10.1101/2024.09.05.611044

**Authors:** Mohini Mendiratta, Meenakshi Mendiratta, Sandeep Rai, Ritu Gupta, Sameer Bakhshi, Mukul Aggarwal, Aditya Kumar Gupta, Hridayesh Prakash, Sujata Mohanty, Ranjit Kumar Sahoo

## Abstract

**Background:** T-cell activation and proliferation are critical for understanding immune responses in both healthy and pathological conditions such as acute graft-versus-host disease (aGVHD). Phytohemagglutinin (PHA) and interleukin-2 (IL-2) are commonly used in in vitro assays to study T-cell responses. In view of the discrete response of T cells from aGvHD patient’s cohorts, our study optimized PHA / IL-2 based T-cell response among healthy individuals versus aGVHD patients.

**Methods:** Peripheral blood was collected from age- and sex-matched healthy individuals (n=10) and aGVHD patients (n=10). CD3^+^ T-cell were isolated and stimulated with varying concentrations of PHA (1-10μg/ml) and IL-2 (50-500 IU/ml). Cell proliferation was assessed using MTS and CFSE assays, while their apoptosis was evaluated with Annexin V/7-AAD staining.

**Results:** We observed enhanced proliferation of healthy individuals at higher PHA concentrations (5-10μg/ml), whereas aGVHD patients exhibited heightened proliferation even at lower PHA concentrations (1-2.5μg/ml) at 48 hours. Prolonged exposure of T cells from GvHD patients to PHA led to decreased proliferation while it increased in the T cells from healthy donors.IL-2 supplementation (50 IU/ml) of T-cells from healthy donors significantly enhanced their proliferation and survival, with the optimal concentration supporting robust proliferation over extended culture periods.

**Conclusion:** Our study optimized PHA and IL-2 concentrations required for T-cell proliferation studies among healthy individuals and aGVHD patients. and underscored experimental conditions required for studying T-cell behavior/dysregulation in aGVHD condition.

**GRAPHICAL ABSTRACT:** 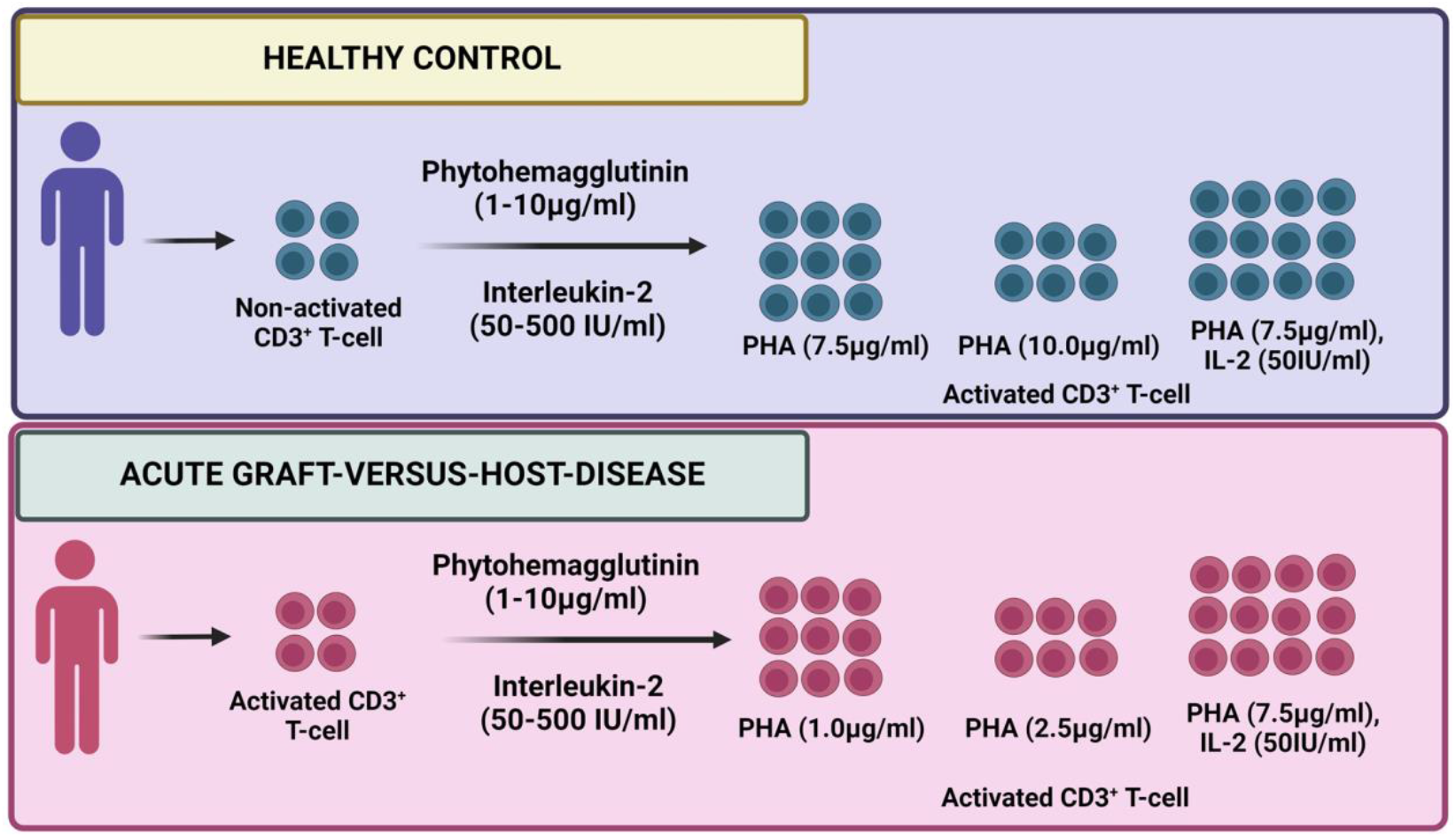

**Highlights of the study:**

- Lower doses of PHA (1.0μg/ml) and IL-2 (50IU/ml) are optimum conditions for aGVHD patients derived CD3+ T-cell proliferation under *in vitro* conditions.
- The maximum T-cell proliferation in healthy individuals occurs with 7.5μg/ml PHA and 50IU/ml IL-2.
- Higher doses of PHA induce cytotoxicity in both cohorts.
- IL-2 significantly enhances T-cell survival, with 50IU/ml maintaining robust proliferation over extended periods.

## 1. Introduction

Phytohemagglutinin (PHA) and interleukin-2 (IL-2) mediated priming/activation of T-cells involves a triggers complex interplay of signaling pathways. This bypasses the normal physiological antigen recognition process which typically requires interaction of TCR with MHC-peptide complexes. [1]. PHA, binds to carbohydrate moieties on the T-cell receptor (TCR) molecules, causing aggregation of these receptors on the cell surface [2]. This triggers the release of calcium (Ca^2+^)-dependent signaling pathways, leading to the translocation of the nuclear factor of activated T-cell (NFAT) into the nucleus [3]. Once in the nucleus, NFAT, often in conjunction with the AP-1 complex, binds to specific DNA sequences, driving the transcription of target genes such as IL-2 [4] which is a critical growth factor for T-cell, is produced in response to PHA-induced NFAT activation. IL-2 binds to its high-affinity receptor on the surface of T-cell, and promotes the transcription of genes necessary for cell proliferation and survival. IL-2 amplifies the T-cell response initiated by PHA by enhancing the proliferation of activated T cells and supporting their survival [1,5]. This synergy between PHA-induced TCR signaling and IL-2-mediated growth signals results in robust T-cell activation and expansion, a process that is crucial for various *in vitro* T-cell-based studies.

Acute Graft-versus-Host-disease (aGVHD) is a post-transplantation autoimmune disorder and one of the most formidable challenges in the allogeneic hematopoietic stem cell transplantation (Allo-HSCT) scenario. In GvHD the graft turns against the host which subsequently leads to a perilous immune battle. It is associated with the failure of our immune system to discriminate among self /nonself which is manifested in autoimmune conditions. where the graft’s immune cells attack the recipient’s tissues [6].

aGVHD is characterized by exaggerated inflammatory responses and hyperactivation of Immune cells and cytokine storm which leads to damage to the recipient’s tissue. This condition is marked by elevated levels of pro-inflammatory cytokines and a dysfunction in regulatory T-cell (Tregs), which encounter the deleterious hyperactive immune response [7].

Excessive T-cell activation and their proliferation is a cardinal sign of aGVHD, which leads to organ damage in patients [9]. Therefore maintaining a balanced T-cell response is paramount for ensuring proper immune functions for imparting immune homeostasis in the host. In this context, setting up T cell-based analysis is inevitable for fine-tuning T cells in aGVHD patient. In this context, Our study optimized T-cell priming regime which are important for both monitoring and modulating T-cell response *in vitro*. Our study demonstrates that Lower doses of PHA (1.0μg/ml) and IL-2 (50IU/ml) are optimum conditions for aGVHD patients-derived CD3+ T-cell proliferation under *in vitro* conditions. The maximum T-cell proliferation in healthy individuals occurs with 7.5μg/ml PHA and 50IU/ml IL-2 which enhances T-cell survival and maintains robust proliferation over extended periods.

We believe that our studies can contribute to optimizing conditions for accessing T-cell functions, dynamics and which can help in designing strategies for repairing T-cell dysregulation in aGVHD.

## 2. Material and methods

### 2.1. Study cohort and ethics approval

Age and sex-matched healthy individuals (n=10) with a median age of 31 years (range 21-41) and aGVHD (grade II-IV) patients (n=10) with a median age of 28.5 years (range 20-43) were recruited prospectively from the Department of Medical Oncology, Dr. B. R. Ambedkar IRCH, Department of Hematology and Pediatrics Oncology, All India Institute of Medical Sciences, New Delhi for the study. The study involved the use of human subjects and was conducted following ethical standards. It was approved by the Institute Ethics Committee for Post Graduate Research, All India Institute of Medical Sciences, New Delhi, India (Ref. No.: IECPG-542/23.09.2020).

Informed written consents were obtained from all participants and all procedures adhered strictly to the guidelines and regulations set forth by the ethics committee.

### 2.2. Isolation and stimulation of CD3^+^ T-cell

Peripheral blood (PB) was collected from healthy individuals and patients in a sterile sodium heparin-coated vacutainer (BD Biosciences, US) and the peripheral blood mononuclear cells (PBMNCs) were isolated using Ficoll density gradient centrifugation according to a standardized protocol [10]. CD3^+^ T-cell was then isolated from PBMNCs by negative selection using a Pan T cell isolation kit, human (Miltenyi Biotec, USA), following the manufacturer’s instructions. Subsequently, CD3^+^ T-cell was stimulated with Phytohemagglutinin (PHA) (Sigma, USA) at concentrations ranging from 1.0μg/ml to 10μg/ml with IL-2 (50IU/ml-500IU/ml) in 1X Rosewell Park’s Memorial Institute (RPMI)-1640 medium supplemented with L-glutamine (Thermo Fisher Scientific, USA), 10% fetal bovine serum (FBS) (Thermo Fisher Scientific, USA), 1% antibiotic-antimycotic solution (Thermo Fisher Scientific, USA) at 37?C, 5% CO_2_ for 24, 48 and 72 hours.

### 2.3. Cell Proliferation using MTS

The proliferation of CD3+ T-cell was assessed using the 3-(4,5-dimethylthiazol-2-yl)-5-(3-carboxymethoxyphenyl)-2-(4-sulfophenyl)-2H-tetrazolium (MTS) (Abcam, US) following the manufacturer’s protocol [11]. Briefly, 10μl of MTS solution was added directly to each well of 96-well plates containing PHA-stimulated and non-activated (control) cells at the specified time points (24, 48, and 72 hours). The plates were then incubated for 3 hours at 37°C. Following incubation, absorbance was measured at 490 nm using an ELISA reader (Bio-Rad, USA), with the blank consisting of culture medium alone. All experiments were conducted with 5 healthy individuals and 5 aGVHD patients, resulting in 5 biological replicates of each. Within each biological replicate, experiments were conducted in triplicates.

### 2.4. CFSE T-cell proliferation assay

CD3^+^ T-cell were labeled with 1μM Cell Trace^TM^ CFSE dye (BD Biosciences, USA) for 20 minutes at 37°C and followed by activation with PHA (1-10μg/ml) (Sigma, USA) in 1X Rosewell Park’s Memorial Institute (RPMI)-1640 medium supplemented with L-glutamine (Thermo Fisher Scientific, USA), 10% FBS, 1% antibiotic-antimycotic solution at 37°C, 5% CO_2_ for 72 hours and 48 hours for healthy subjects and aGVHD patients respectively. Additionally, the cell proliferation was assessed with PHA and IL-2 (50IU/ml) using CFSE T-cell proliferation assay [12].

All experiments were conducted with 10 biological replicates of each healthy and aGVHD cohort.

### 2.5. Apoptosis assay

Upon stimulation of T-cell, the cell suspensions were collected, washed, and stained with Annexin V/7AAD (BD Pharmingen^TM^, USA) using the manufacturer’s instructions [13]. The cells were acquired using a DxFlex flow cytometer (Beckmann Coulter) for the enumeration of viable, apoptotic, and necrotic cells, and the data was analyzed using Kaluza software version 2.1 (Beckmann Coulter).

### 2.6. Statistical analysis

All statistical analyses were conducted using GraphPad Prism version 8.4.3. One-way and Tukey’s post hoc tests compared three or more groups. Data was shown as Mean±S.D. and a p-value of ≤ 0.05 was considered statistically significant.

## 3. Results

### 3.1. Temporal variations in PHA-driven CD3+ T-cell proliferation in healthy and aGVHD patients

Initially, we evaluated the proliferation of CD3^+^ T-cell derived from healthy individuals and aGVHD patients using the MTS assay at various concentrations of PHA (1-10μg/ml) at various time points (24, 48 and 72 hours). We observed that there was no significant difference in the proliferation of activated CD3^+^ T-cell derived from both healthy individuals and aGVHD compared to their respective non-activated controls at 24 hours, irrespective of PHA concentration (Figure 1A).

**Figure 1:**
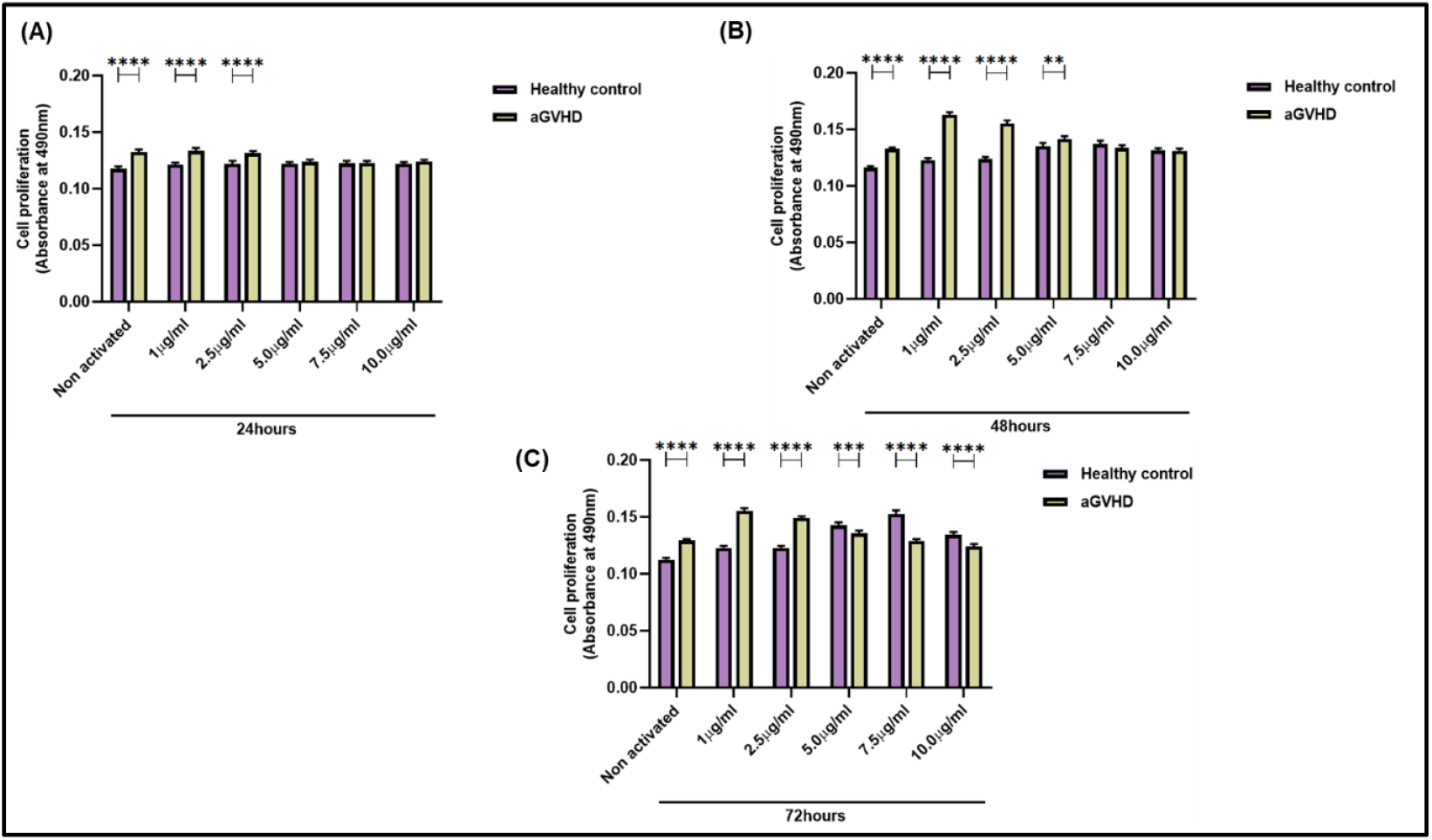
CD3^+^ T cell proliferation of healthy control and aGVHD patients with and without different doses of PHA (1μg/ml-10μg/ml) at various time points using MTS assay. The bar graph represents proliferation (a) 24 hours, (B) 48 hours, and (C) 72 hours.

At 48 hours, there was a noticeable increase in cell proliferation in healthy individuals, particularly at PHA concentrations of 5-10μg/ml. In contrast, aGVHD patients exhibited increased proliferation across all PHA concentrations (1-5μg/ml) from their non-activated ones, with a major increase at 1 and 2.5μg/ml, followed by a decline at higher concentrations, although still higher than at 24 hours (Figure 1B). At 72 hours, a slight decrease in proliferation was observed in CD3+ T-cell derived from aGVHD patients compared to 48 hours at all PHA concentrations (Figure 1C). In healthy individuals, however, there was a significant increase in proliferation at 72 hours, with the highest proliferation observed at PHA concentrations of 5.0μg/ml and 7.5μg/ml (Figure 1C). Overall, we observed that higher PHA concentration led to a decrease in CD3+ T-cell proliferation in both healthy individuals and aGVHD patients. Moreover, prolonged exposure to PHA further reduced the proliferation of CD3+ T-cell derived from aGVHD patients.

### 3.2. Differential CD3^+^ T-cell proliferation and activation status in healthy individuals and aGVHD patients

Our findings indicated that the maximum proliferation of CD3^+^ T-cell occurred at 72 hours for healthy individuals and at 48 hours for aGVHD patients. This observation was initially inferred through the MTS assay, which primarily measures metabolic activity as an indirect indicator of cell proliferation. To directly observe cell proliferation, we employed the CFSE proliferation assay at 72 hours and 48 hours for healthy individuals and aGVHD patients, respectively, across all PHA concentrations (1-10μg/ml).

Morphological analysis revealed a notable variation in CD3^+^ T cell morphology following stimulation. Clumping of cells was evident in PHA-stimulated CD3+ T cells, particularly in aGVHD patients, where clumps were observed even in non-activated cells. Interestingly, small clumps were present at lower PHA concentrations (1.0μg/ml) in aGVHD patient-derived cells, with increasing PHA concentrations leading to more pronounced clumping and a rise in necrotic cells within these clumps (Figure 2). In healthy individuals, cell clumping was primarily observed at higher PHA concentrations (5.0-10μg/ml), with a slight reduction in clumping at 10μg/ml compared to 5.0 and 7.5μg/ml (Figure 2). These observations suggest that higher PHA concentrations may exert toxic effects on the cells and this clumping is due to increased expression of adhesion molecules on the cell surface, facilitating stronger cell-cell interactions.

**Figure 2:**
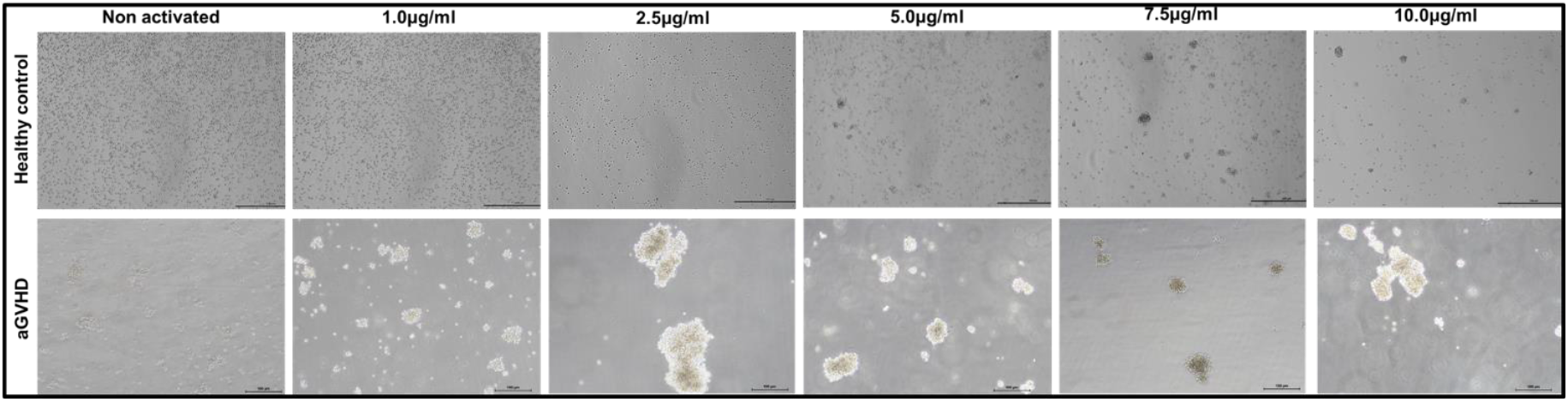
Representative morphological images of CD3+T cell of healthy control and aGVHD patients with and without different doses of PHA (1.0μg/ml-10μg/ml) at 72 hours and 48 hours respectively.

Further analysis using the CFSE assay confirmed these proliferation patterns. In healthy individuals, we observed there was a significant increase in the proliferation at PHA concentrations of 5.0-10μg/ml, with a maximum of 7.5μg/ml from non-activated ones (94.772% vs. 0.0286%; p ≤ 0.0001) (Figure 3A). In contrast, aGVHD patient-derived cells exhibited increased proliferation at lower PHA concentrations (1.0-2.5 μg/ml), followed by a decline in proliferation at higher concentrations (5.0-10μg/ml). Interestingly, the maximum proliferation of aGVHD patients-derived CD3^+^ T-cell was observed at 1.0μg/ml compared to 2.5μg/ml (93.948% vs. 90.246%; p ≤ 0.0001) (Figure 3A). This pattern suggests that the CD3^+^ T-cell from aGVHD patients were pre-activated, likely due to the inflammatory environment associated with the disease.

**Figure 3:**
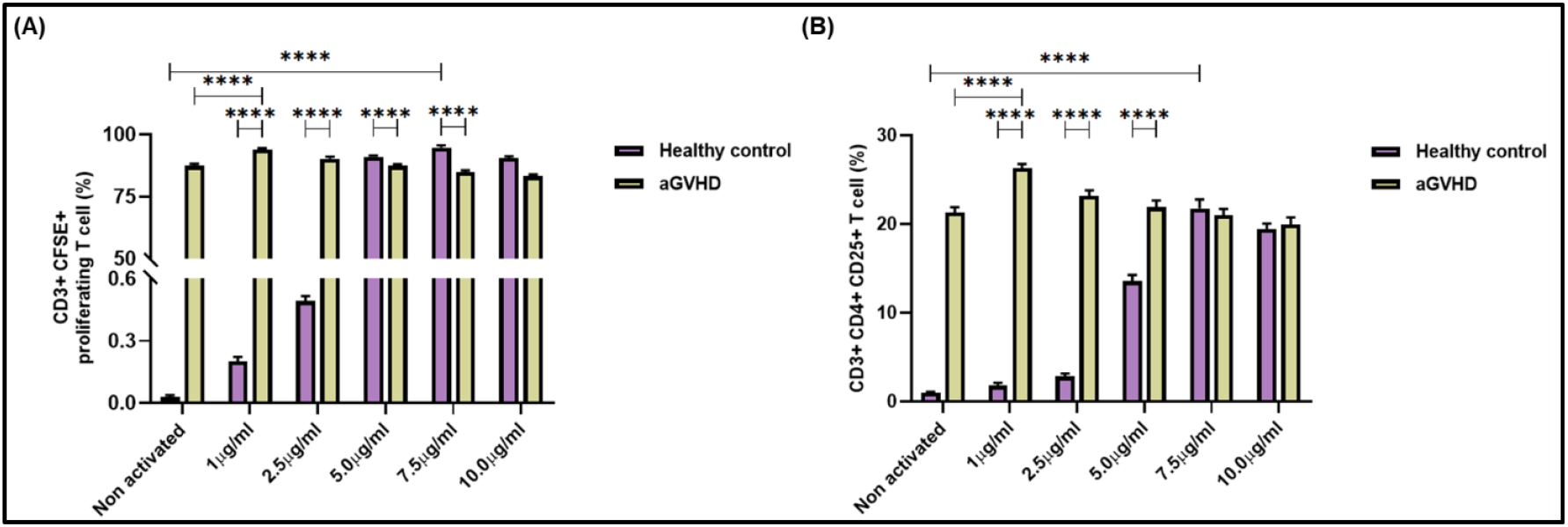
CD3^+^ T cell proliferation of healthy control (n=5) and aGVHD patients (n=5) at 72 hours and 48 hours respectively with and without different doses of PHA (1μg/ml-10μg/ml) using flow cytometry. The bar graph represents (A) CD3^+^ CFSE^+^ proliferating T cell (%) using Cell Trace^TM^ CFSE cell proliferation assay, (B) CD3^+^ CD4^+^ CD25^+^ T cell (%) using flow cytometry.

To further confirm the activation status of these cells, we evaluated the expression of CD25 on helper T-cell. In aGVHD patients, there was an increased expression of CD25 at lower PHA concentrations, with the highest expression observed at 1.0 μg/ml compared to non-activated ones (21.316% vs 26.332%; p ≤ 0.0001) (Figure 3B). Conversely, in healthy individuals, CD25 expression was enhanced at higher PHA concentrations (5.0-10μg/ml), with the most notable expression at 7.5μg/ml compared to non-activated ones (21.77% vs. 0.994%; p ≤ 0.0001) (Figure 3B). These findings align with the CFSE-based cell proliferation assay, corroborating the differential response of CD3+ T cells to PHA stimulation between healthy individuals and aGVHD patients.

### 3.3. Higher doses of PHA cause cytotoxicity of CD3^+^ T-cell in healthy individuals and aGVHD patients

To determine the optimum dose of PHA that promotes healthy T-cell proliferation without inducing cytotoxic effects, we performed an Annexin V/7-AAD staining assay. This was crucial to assess the viability of CD3^+^ T-cell at various PHA concentrations, particularly in healthy individuals and aGVHD patients.

Our analysis revealed a significant difference in the proliferation response to PHA between healthy individuals and aGVHD patients. In healthy individuals, maximum proliferation was observed at PHA concentrations of 7.5μg/ml and 10.0μg/ml after 72 hours of exposure. However, as the PHA concentration increased, a corresponding increase in cytotoxicity was noted. Specifically, the proportion of viable cells decreased from 93.45% to 90.238% (p ≤ 0.0001) at 7.5μg/ml and 10.0μg/ml respectively, indicating that higher PHA concentrations might exert a cytotoxic effect on T-cell despite their enhanced proliferation (Figure 4A, B).

**Figure 4:**
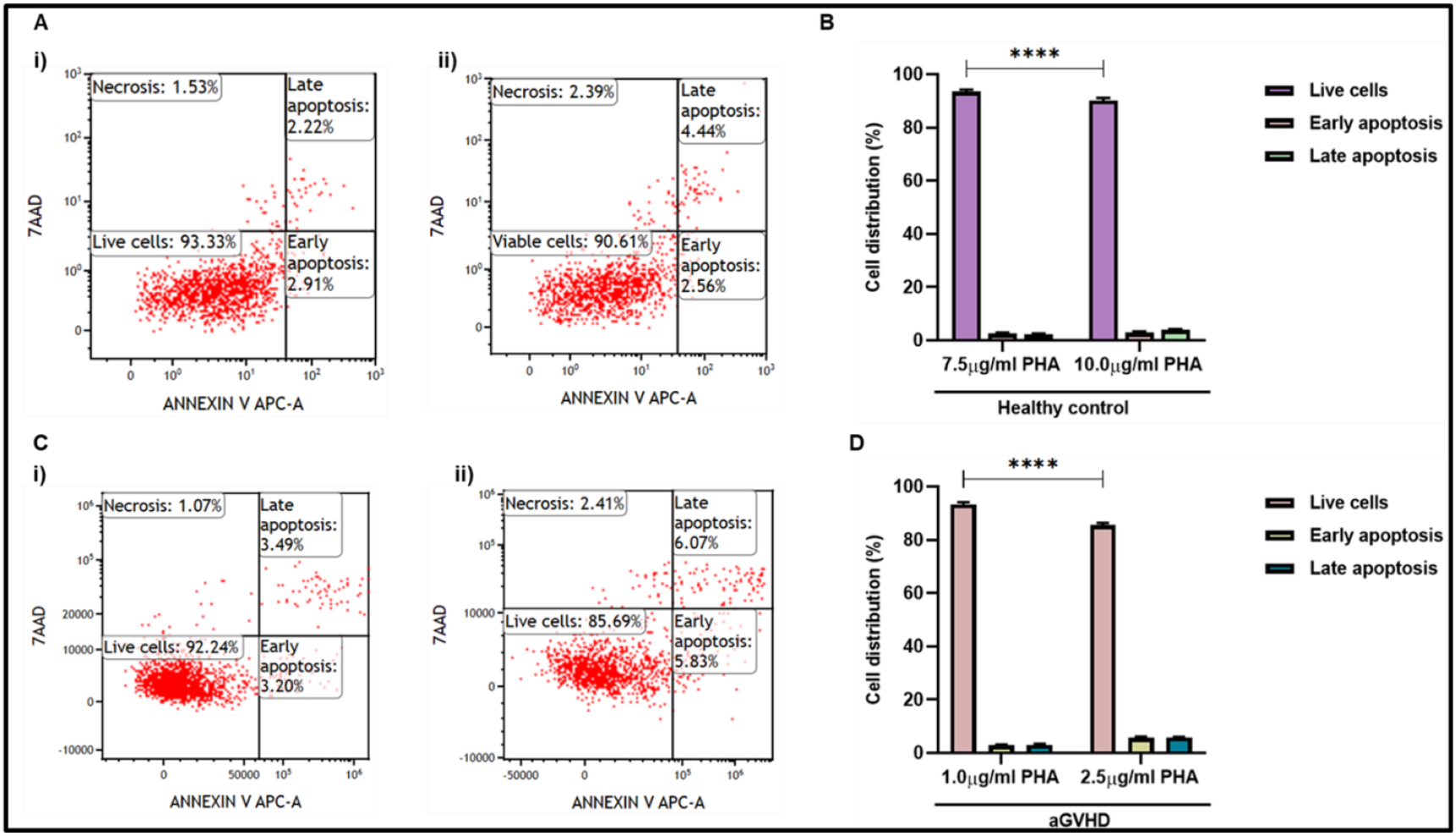
Cell distribution of CD3^+^ T cell of healthy control (n=5) and aGVHD (n=5) patients using Annexin-V/7AAD staining. Representative dot plots of cell distribution of (A) Healthy control at different PHA concentrations i) 7.5μg/ml, ii) 10.0μg/ml; (C) GVHD patients at different PHA concentrations i) 1.0μg/ml, ii) 2.5μg/ml. The bar graphs represent cell distribution in percentage of (B) Healthy control, and (D) aGVHD patients at different PHA concentrations.

In contrast, we observed a significant increase in the proliferation of aGVHD-derived CD3^+^ T-cell at lower PHA concentrations (1.0 and 2.5μg/ml). The viability of these cells was notably higher at 1.0μg/ml than 2.5μg/ml (93.332% vs. 85.436%; p ≤ 0.0001) (Figure 4C, D). This suggests that even modest increases in PHA concentration can significantly impact the survival of T-cell in aGVHD patients.

Overall, our findings underscore the importance of selecting an appropriate PHA concentration to maximize T-cell proliferation while minimizing cytotoxic effects. For healthy individuals, a concentration around 7.5μg/ml appears optimal, whereas, for aGVHD patients, a lower concentration of 1.0μg/ml is preferable to avoid cytotoxicity and ensure a healthy proliferation response. These results highlight the differential sensitivity of T-cell from healthy individuals versus aGVHD patients to PHA, reinforcing the need for tailored approaches in experimental and therapeutic settings.

### 3.4. Impact of IL-2 on T-cell proliferation and survival in healthy individuals and aGVHD patients

We evaluated the effect of IL-2 across a range of concentrations (50IU/ml to 250IU/ml) on CD3^+^ T-cell derived from healthy individuals and aGVHD patients to optimize the *in vitro* conditions for T-cell survival and proliferation using MTS assay. We monitored cell proliferation with 7.5μg/ml and 1μg/ml of PHA for healthy individuals and aGVHD patients respectively from day 3 to day 10.

Our findings demonstrated a marked increase in T-cell proliferation in the presence of IL-2 compared to conditions without IL-2 across both study cohorts. Notably, during the initial phase (day 3 to day 5), we observed an enhanced proliferation at lower IL-2 concentrations (50IU/ml to 150IU/ml). Specifically, the proliferation peaked at 50IU/ml and 100IU/ml, indicating that these concentrations effectively support T-cell growth during the early stages of culture (Figure 5A, B).

**Figure 5:**
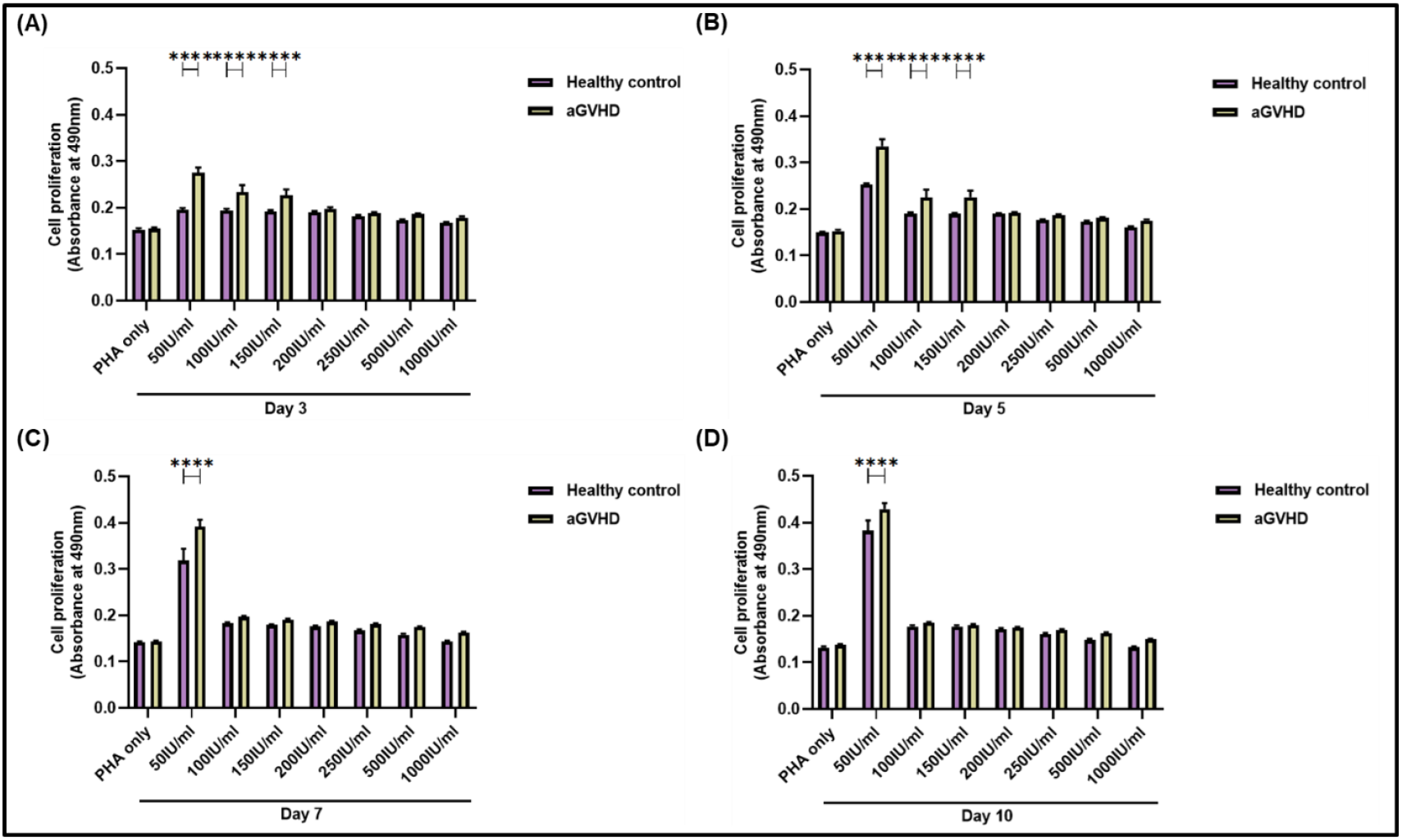
CD3^+^ T cell proliferation of healthy control (n=5; PHA=7.5μg/ml) and aGVHD (n=5; PHA=1.0 μg/ml) patients with and without different doses of IL-2 (50IU/ml-1000IU/ml) at different time points using MTS assay. The bar graph represents proliferation (A) Day 3, (B) Day 5, (C) Day 7, and (D) Day 10.

However, as the culture period was extended to day 7 and day 10, a decrease in proliferation was noted at IL-2 concentrations of 100IU/ml and 150IU/ml in both healthy individuals and aGVHD patients (Figure 5C, D). This decline suggests that higher IL-2 concentrations may become less effective or potentially induce a level of IL-2-driven exhaustion or cytotoxicity over prolonged periods.

Overall, the data suggest that 50IU/ml of IL-2 consistently supports robust T-cell proliferation from day 3 to day 10 in both healthy individuals and aGVHD patients. This concentration was found to be the most effective for maintaining cell viability and proliferation over the extended culture period, highlighting its potential as an optimal dose for *in vitro* T-cell studies.

### 3.5. Optimized PHA and IL-2 conditions drive robust T-cell proliferation in healthy individuals and aGVHD patients

We conducted an *in vitro* proliferation assay on CD3+ T-cell derived from healthy individuals and aGVHD patients under their respective conditions of PHA (7.5μg/ml and 1.0μg/ml) and IL-2 (50IU/ml) using the CFSE proliferation. Our findings demonstrated that 94.33% of CD3+ T-cell in healthy individuals and 93.8% in aGVHD patients were actively proliferating under these conditions (Figure 6A, B, C). This finding underscores the effectiveness of the selected PHA concentrations in conjunction with IL-2 in driving T-cell proliferation. The ability of these conditions to maintain high proliferation rates highlights their potential utility in further studies aimed at understanding T-cell behavior and immune modulation in both healthy and disease states. Additionally, the slight difference in proliferation between the two cohorts reflects the tailored approach needed to optimize T-cell activation based on specific patient profiles.

**Figure 6:**
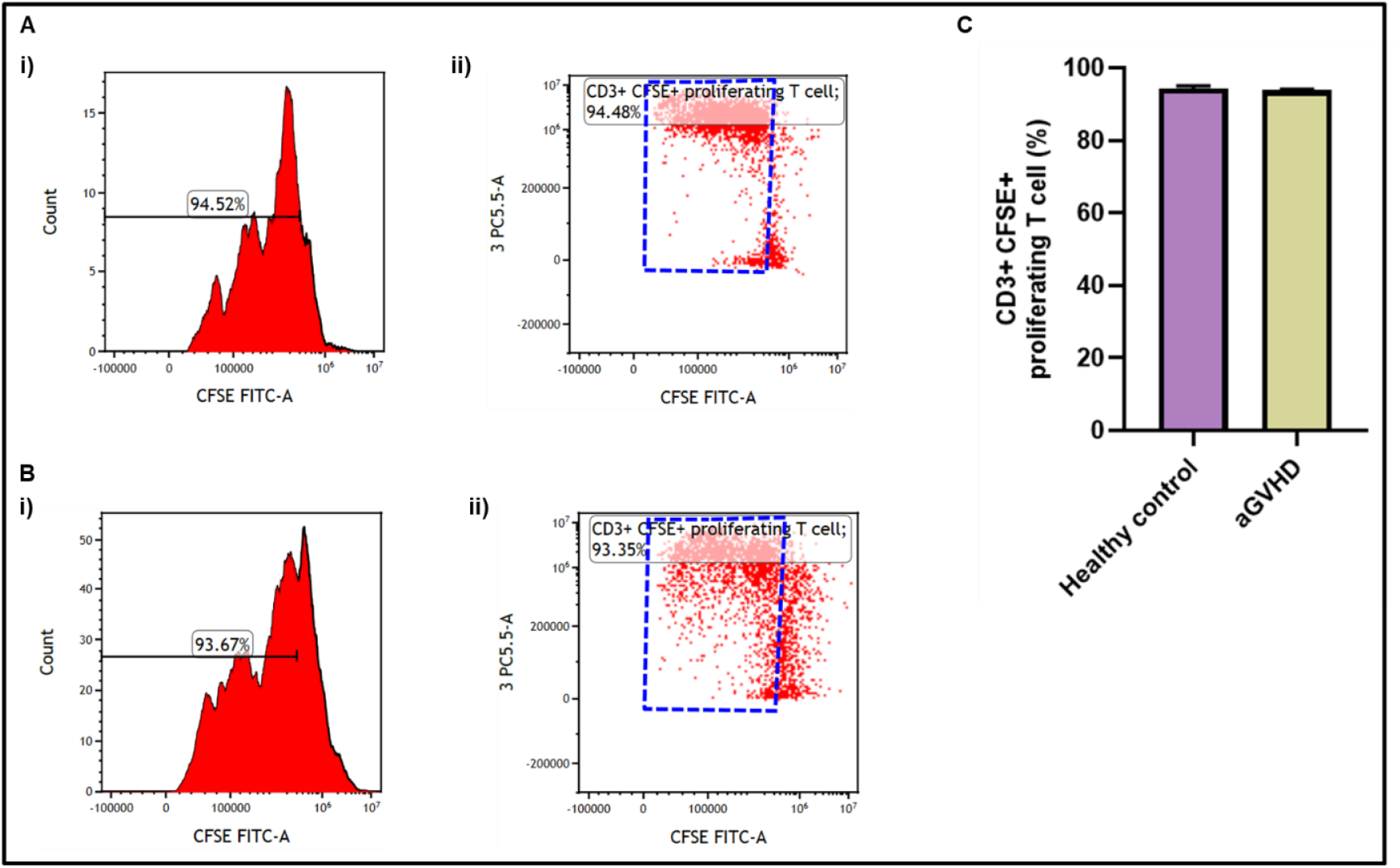
CD3^+^ T cell proliferation of healthy control (n=5) and aGVHD (n=5) patients using CellTrace^TM^ CFSE cell proliferation assay. Representative histograms (i) and dot plots (ii) of T cell proliferation of (A) Healthy control (7.5μg/ml; 50IU/ml) and (B) aGVHD patients (1.0μg/ml; 50IU/ml). (C) The bar graphs represent the CD3+ CFSE+ proliferating T cell percentage in healthy control and aGVHD.

## 4. Discussion

Our study provides new insights into the differential responses of CD3^+^ T-cell from healthy individuals and aGVHD patients to PHA stimulation /IL-2 supplementation. Our findings reveal distinct variations in proliferation, activation, and cytotoxicity in the T cells which emphasizes the need for tailored experimental conditions or maintaining optimum response in T cells.

Our results showed that CD3+ T-cells from healthy individuals exhibited increased proliferation at 48 and 72 hours in response to PHA, particularly at concentrations of 5-10.0 μg/ml, with the most significant increase observed at7.5 μg/ml which corroborates previous studies indicating that optimal T-cell proliferation in healthy individuals requires a moderate level of stimulation. A study demonstrated that T-cell respond to PHA in a dose-dependent manner, suggesting that T cells become tolerant to excessive stimulation [14]. Moreover, it was reported that PHA can be a good alternative to CD3/CD28 beads [15] for efficient T-cell proliferation with reduced exhaustion markers T cells from aGVHD patients exhibit heightened proliferation even at lower PHA concentrations (1-2.5 μg/ml) indicating their hypersensitivity which could be due to elevated levels of NFAT/ PKC and calcium pathways which impart them a constitutive pre-activated phenotype in aGVHD [16].

Morphological changes, such as cell clumping, were more pronounced in aGVHD patient-derived T-cells even at lower PHA concentrations. This clumping is indicative of increased cell-cell adhesion, possibly due to the increased expression of adhesion molecules such as CD54, and integrin/cadherin / CLA etc. The presence of necrotic cells within these clumps could be due to high ROS and/or Th1/ 17 responses, particularly at higher concentrations. These findings demonstrate differential sensitivity of T-cells from healthy / aGvHD patients to PHA stimulation. Previous studies have demonstrated that T-cell undergo morphological changes upon activation, transitioning from single, spherical cells to clumped formations. This change is triggered by the activation of the T-cell receptor (TCR), which leads to Ca2+ mobilization [17–19].

Our cytotoxicity assays revealed that PHA stimulation (irrespective of concentration) induced more toxicity in the T-cell pool from aGVHD patients which could be due to ROS and Th17 programming of T-cell [15,20–22].

Transient exposure of T-cell populations both healthy and aGvHD patients to IL-2 significantly enhanced their proliferation. It aligns with previous research demonstrating the critical role of IL-2 in sustaining T-cell proliferation and preventing apoptosis [23]. However, longer exposure of these cells to IL-2 diminished their proliferation which could be due to IL-2-driven exhaustion or cytotoxicity. It is consistent with previous observations that prolonged IL-2 exposure can lead to T-cell exhaustion [24] and a lower dose of IL-2 is more effective in T-cell proliferation and survival under *in vitro* conditions [25]. The data suggest that a concentration of 50 IU/ml IL-2 is optimal for maintaining robust T-cell proliferation and viability over time.

Our optimized conditions of PHA and IL-2 demonstrated high levels of T-cell proliferation in both healthy and aGVHD patient samples. Similar approaches have been validated in other studies that emphasize the importance of optimizing stimulation conditions for effective T-cell proliferation in various clinical and research settings [1,26]. This underscores the effectiveness of these conditions in driving robust T-cell proliferation and highlights their potential utility in further studies of T-cell immunological behavior.

Although our study focused on the effects of PHA and IL-2 on T-cell proliferation and activation, however, the influence of other factors influencing T-cell behavior, such as cytokine milieu and interactions with other immune cells need further investigations which would provide a more comprehensive understanding of T-cell dynamics in aGVHD and other immune-mediated conditions.

## 5. Conclusion

In summary, our study provides valuable insights into the differential responses of T-cells to various cognitive mitogens among healthy individuals and aGVHD patients. The study discuss the importance of optimizing experimental conditions for T-cell activation and sensitivity variations which is important for designing new immune therapeutics for aGvHD condition.

## Funding acknowledgement

The study has been supported by the Indian Council of Medical Research, New Delhi, India (Grant Id: 2021/14763).

## 6. Credit authorship contribution statement

**Mohini Mendiratta:** Methodology, Data curation, Formal analysis, Writing original draft, Review, and Editing; **Meenakshi Mendiratta:** Methodology, Data curation and Formal analysis; **Sandeep Rai:** Data curation and Formal analysis; **Ritu Gupta:** Resources and Data analysis; **Sameer Bakhshi:** Resources; **Mukul Aggarwal:** Resources; **Aditya Kumar Gupta:** Resources; **Hridayesh Prakash:** Conceptualization, Supervision, Data curation, Formal analysis, Review, and Editing; **Sujata Mohanty:** Conceptualization, Resources, Supervision, Data curation, Formal analysis, Review, and Editing; **Ranjit Kumar Sahoo:** Conceptualization, Methodology, Supervision, Funding, Resources, Data curation, Formal analysis, Review, and Editing.

## 8. Data availability

The data that support the findings of this study will be made available from the corresponding author upon reasonable request.

## 9. Acknowledgements

The authors express their gratitude to the All India Institute of Medical Sciences (AIIMS), New Delhi, India for facilitating the execution of the study. A graphical abstract was created using Biorender.com.

## 10. Abbreviations

PHA: Phytohemagglutinin
IL-2: Interleukin-2
TCR: T-cell receptor
NFAT: Nuclear Factor of Activated T-cell
aGVHD: Acute Graft-versus-Host-Disease
Tregs: Regulatory T-cell
MTS: 3-(4,5-dimethylthiazol-2-yl)-5-(3-carboxymethoxyphenyl)-2-(4-sulfophenyl)-2H-tetrazolium
CFSE: carboxyfluorescein succinimidyl ester
7AAD: 7-amino actinomycin D

